# Genome-wide characterization of *Phytophthora infestans* metabolism: a systems biology approach

**DOI:** 10.1101/171082

**Authors:** Sander Y.A. Rodenburg, Michael F. Seidl, Dick de Ridder, Francine Govers

## Abstract

Genome-scale metabolic models (GEMs) provide a functional view of the complex network of biochemical reactions in the living cell. Initially mainly applied to reconstruct the metabolism of model organisms, the availability of increasingly sophisticated reconstruction methods and more extensive biochemical databases now make it possible to reconstruct GEMs for less characterized organisms as well, and have the potential to unravel the metabolism in pathogen-host systems. Here we present a GEM for the oomycete plant pathogen *Phytophthora infestans* as a first step towards an integrative model with its host. We predict the biochemical reactions in different cellular compartments and investigate the gene-protein-reaction associations in this model to get an impression of the biochemical capabilities of *P. infestans.* Furthermore, we generate life stage-specific models to place the transcriptomic changes of genes encoding metabolic enzymes into a functional context. In sporangia and zoospores there is an overall downregulation, most strikingly reflected in the fatty acid biosynthesis pathway. To investigate the robustness of the GEM, we simulate gene deletions to predict which enzymes are essential for *in vitro* growth. While there is room for improvement, this first model is an essential step towards an understanding of *P. infestans* and its interactions with plants as a system, which will help to formulate new hypotheses on infection mechanisms and disease prevention.

## Introduction

The growth and functioning of any living cell is governed by a complex, interconnected set of biochemical reactions, comprehensively referred to as its metabolism (Nielsen, 2017). It is essential for cells to consume and break down nutrients taken from the environment, and to use the resulting basic building blocks to construct the molecules needed for life (nucleic acids, amino acids, lipids etc.) and for survival (secondary metabolites). However, the many molecules in this system and the many parameters that govern the biochemical reactions make metabolism hard to study. Systems biology was introduced as a method to study a biological system as a whole by capturing its behaviour in a mathematical abstraction, i.e. a model (Ideker *et al.*, 2001). A model can provide insights into the response of a biological system to certain perturbations or stimuli (Bordbar *et al.*, 2014). A widely studied class of models is that of genome-scale metabolic models (GEMs), which simulate and predict the metabolic behaviour of a cell (Lewis *et al.*, 2012), such as the nutrients it can assimilate and the molecules it can synthesize.

The foundation of a GEM is the set of biochemical reactions that may occur in a cell, often catalyzed by enzymes. Hence, the identification of enzyme encoding genes in the genome of an organism can help to reconstruct an overview of its biochemical capabilities (O’Brien *et al.*, 2015; Yilmaz and Walhout, 2016). In a metabolic model, every reaction is considered as a conversion of substrate metabolites into product metabolites that takes place at a specific rate. The stoichiometry represents the balance of metabolites within the reaction. In steady state, i.e. a situation in which the net metabolite concentrations do not change, the reaction rates are called fluxes. A class of methods called constraint-based modelling can be used to simulate the distribution of these fluxes in certain conditions (Orth *et al.*, 2010). A well-known constraint-based method is flux balance analysis (FBA), which calculates the optimal set of flux values for the entire GEM to attain a specific metabolic objective. Typically, this metabolic objective is maximization of biomass production, a synonym for growth, but can also entail different objectives, for instance, minimization of energy consumption or redox potential (García Sánchez and Torres Sáez, 2014).

To date several semi-automated GEM reconstruction methods and protocols have been proposed (Agren *et al.*, 2013; Karp *et al.*, 2009; Schellenberger *et al.*, 2011; Thiele *et al.*, 2014; Thiele and Palsson, 2010), and the development of central databases for metabolic pathways and models has made biochemical information widely available (Caspi *et al.*, 2014; Kanehisa *et al.*, 2015; King *et al.*, 2016). While initially GEM reconstruction was mainly limited to microbes (prokaryotes and simple eukaryotes), the available resources now allow for reconstruction of genome-scale metabolic models for complex organisms such as mammals and higher plants (Dharmawardhana *et al.*, 2013; Thiele *et al.*, 2013; Yuan *et al.*, 2016). Such models have also already been applied to understand the metabolic interactions between pathogen and host (Duan *et al.*, 2013; Huthmacher *et al.*, 2010; Peyraud *et al.*, 2016). This can provide new hypotheses about a pathogen’s infection strategy and may suggest novel control targets (Chavali *et al.*, 2011; Sharma *et al.*, 2017).

*Phytophthora infestans* is the causal agent for the devastating late blight disease on tomato and potato, posing an important threat to global food production. It is considered one of the model species in the oomycete taxonomic lineage that comprises a miscellaneous collection of eukaryotes, many of which are pathogenic on plants or animals (Haas *et al.*, 2009). In the asexual life cycle of *P. infestans* different stages can be distinguished (Judelson, 2017). When the mycelium starts to sporulate it forms sporangia that are dispersed by wind and water. Sporangia either germinate directly starting new infections or develop into zoosporangia that release zoospores. The latter encyst upon plant contact and germinate, thereby forming an appressorium at the tip from which a penetration peg emerges that mediates entry into the epidermal cells of the host plant. Cell wall degrading enzymes are secreted that may facilitate the penetration process (Brouwer *et al.*, 2014; Meijer *et al.*, 2014). After penetration, hyphae colonize the mesophyll where they grow intracellular and form haustoria inside the host cells (Whisson *et al.*, 2016). These feeding structures provide a large contact area with the host cytosol, enabling efficient exchange of molecules, to mediate further infection. Apart from the pathogen-host interactions at the protein level, it can be anticipated that an unknown combination of metabolites is taken up from the plant by the pathogen as nutrients.

*P. infestans* is able to assimilate a wide range of compounds (Hohl, 1991). *In vitro*, *P. infestans* is for example able to grow on pea, rye or Henninger medium which contains an undetermined mixture of various nutrients, such as amino acids and organic acids and lipids (Griffiths *et al.*, 2003; Meijer *et al.*, 2014). Many of the *Peronosporales*, the lineage that comprises the *Phytophthora* genus, are sterol and thiamine auxotroph, which implies that these compounds must be acquired from the host (Dahlin *et al.*, 2017; Gaulin *et al.*, 2010; Judelson, 2012); while sterols are not essential (yet highly beneficial) for mycelial growth, thiamine is (Hohl, 1991). The nutrients that are taken up by the pathogen are converted into biomass and secondary metabolites. *P. infestans* forms various long-chain polyunsaturated fatty acids, predominantly arachidonic- and eicosapentaenoic acid (Griffiths *et al.*, 2003; Sun *et al.*, 2013). The oomycete cell wall is composed of various sugar polymers, mainly 1,3- and 1,6-β-glucans and cellulose (Grenville-Briggs *et al.*, 2008). Notably, both the long-chain polyunsaturated fatty acids and the cell wall glucans can elicit plant immune responses (Robinson and Bostock, 2015), but it is likely that during infection such responses are suppressed by secreted effector proteins.

Large transcriptional changes of genes encoding metabolic enzymes were observed during the asexual lifecycle of *P. infestans* (Ah-Fong *et al.*, 2017), suggesting profound changes at the metabolic level. Notably, metabolic enzymes in general were downregulated in the sporangia and zoospores, and many metabolic processes (e.g. biosynthesis of various amino acids) were upregulated in cysts, and during mycelial growth (Ah-Fong *et al.*, 2017; Grenville-Briggs *et al.*, 2005). Moreover, elevated expression *in planta* of various nutrient transporter genes suggest a rich influx of nutrients during infection (Abrahamian *et al.*, 2016). Transcriptome studies have analysed the metabolism of *P. infestans* from a regulatory point of view. However, these studies do not consider post-transcriptional regulation and metabolic reaction fluxes. A genome-scale metabolic model can provide an overview of the *P. infestans* metabolism, and at the same time predict the functioning of the primary metabolism as a system. Here we propose a first genome-scale metabolic model for *P. infestans.*

## Results & Discussion

### Draft model reconstruction

We identified the enzymes encoded in the *Phytophthora infestans* genome (Haas *et al.*, 2009) by matching all predicted protein sequences to hidden Markov models (HMMs), trained on groups of orthologous proteins from the KEGG Orthology (KO) database (Agren *et al.*, 2013; Kanehisa *et al.*, 2015). This is a particularly suitable method for detection of distant orthologs, since conserved domains have a strong influence on the alignment score and thus this method is sensitive to conserved catalytic domains (Pearson, 2013). Roughly 32% (5,856) of the 18,140 predicted *P. infestans* proteins matched a KO group, yet not every KO group represents a metabolic enzyme catalyzing a biochemical reaction. In total, 1,408 *P. infestans* genes were associated with 1,569 different biochemical reactions, involving 1,663 different metabolites.

*P. infestans* is able to assimilate a range of nitrogen compounds, preferably amino acids but also inorganic forms such as nitrate (Hohl, 1991). As a carbon source, *P. infestans* prefers glucose or sucrose, but can also utilize many mono- and disaccharides (Judelson, 2017). Early experiments determined that *P. infestans* can utilize a range of organic sulphur and phosphor compounds, although more optimal growth rates were observed for inorganic sulphate and phosphate sources (Fothergill and Child, 1964). We added uptake reactions to the model for the minimal synthetic growth medium from literature (Hohl, 1991), the simplest nutrient combination shown to yield *in vitro* growth: glucose, ammonia, phosphate, sulphate and thiamine. Next, we composed a pool of biomass precursor metabolites that must be produced to sustain life: all nucleotides, all 20 L-type amino acids, energy carriers (ATP, GTP), and the cofactors Coenzyme-A, NADH, NADPH and FADH_2_, that are generally essential for a eukaryotic cell (Nielsen, 2017). The exact relative abundance of biomass components has never been quantified for *P. infestans*, therefore the aforementioned biomass metabolites were added to the model as substrates of a single artificial biomass reaction with equal stoichiometry. Additionally, for the phospholipids and fatty acids detected in *P. infestans* (Griffiths *et al.*, 2003) excretion reactions were included. The known cell wall components 1,3- and 1,6- β-glucan and cellulose are all polysaccharides for which glucose is the precursor metabolite.

We used flux balance analysis (FBA) to calculate the flux through each reaction, optimizing for biomass production (Orth *et al.*, 2010). To predict quantitative fluxes using FBA, it is required to provide: an accurate biomass composition, maintenance ATP requirements, growth rates and species-specific reaction constraints (Thiele and Palsson, 2010). While the lack of detailed data on *P. infestans* metabolism currently impairs reliable quantitative flux predictions, we can nevertheless deploy FBA to interrogate the model for its connectivity and topology.

Metabolic enzymes are located in various organelles, causing specific metabolic processes to take place in different parts of the cell. For example, the TCA cycle is typically found in the mitochondria (Zimorski *et al.*, 2017). There is an extensive exchange of metabolites between subcellular compartments (Wanders *et al.*, 2016). Obviously, the compartmentalization influences the connectivity of the reactions and the global behaviour of the model. Based on localization predictions by LocTree 3 (Goldberg *et al.*, 2014), we expanded the model by dividing *P. infestans* proteins over seven subcellular compartments (Figure 1). Reactions in the model were assigned to a particular compartment if at least one of the associated enzymes was predicted to localize there. LocTree has been trained on general eukaryotic sequences, which could influence the accuracy of our enzyme localization predictions. However, previous analyses using similar localization predictors showed that proteins predicted to co-localize are often also co-expressed in *P. infestans* (Seidl *et al.*, 2013). The cytosol contained 1,119 reactions while the mitochondria contained 359, which is approximately 15% of the total number of reactions in the model (Table 1). Of these 359, 160 (45%) were shared with the cytosol (Supplementary Figure S1). Notably, these shared reactions are part of various metabolic pathways, but with a relatively large number (42) in the fatty acid biosynthesis (FAB) pathway. The elongation of fatty acids can be governed by a single fatty acid synthase enzyme (EC 2.3.1.86). *P. infestans* has three gene copies for this enzyme, one of which is predicted to encode a mitochondrial isoform (PITG_18025). It was reported that many eukaryotes have a highly conserved, independent mitochondrial FAB pathway that is crucial for development (Hiltunen *et al.*, 2009; Kastaniotis *et al.*, 2017). *Phytophthora* spp. are thought to store energy in fatty acid molecules to facilitate movement of zoospore flagellae (Judelson, 2017).

**Figure 1:**
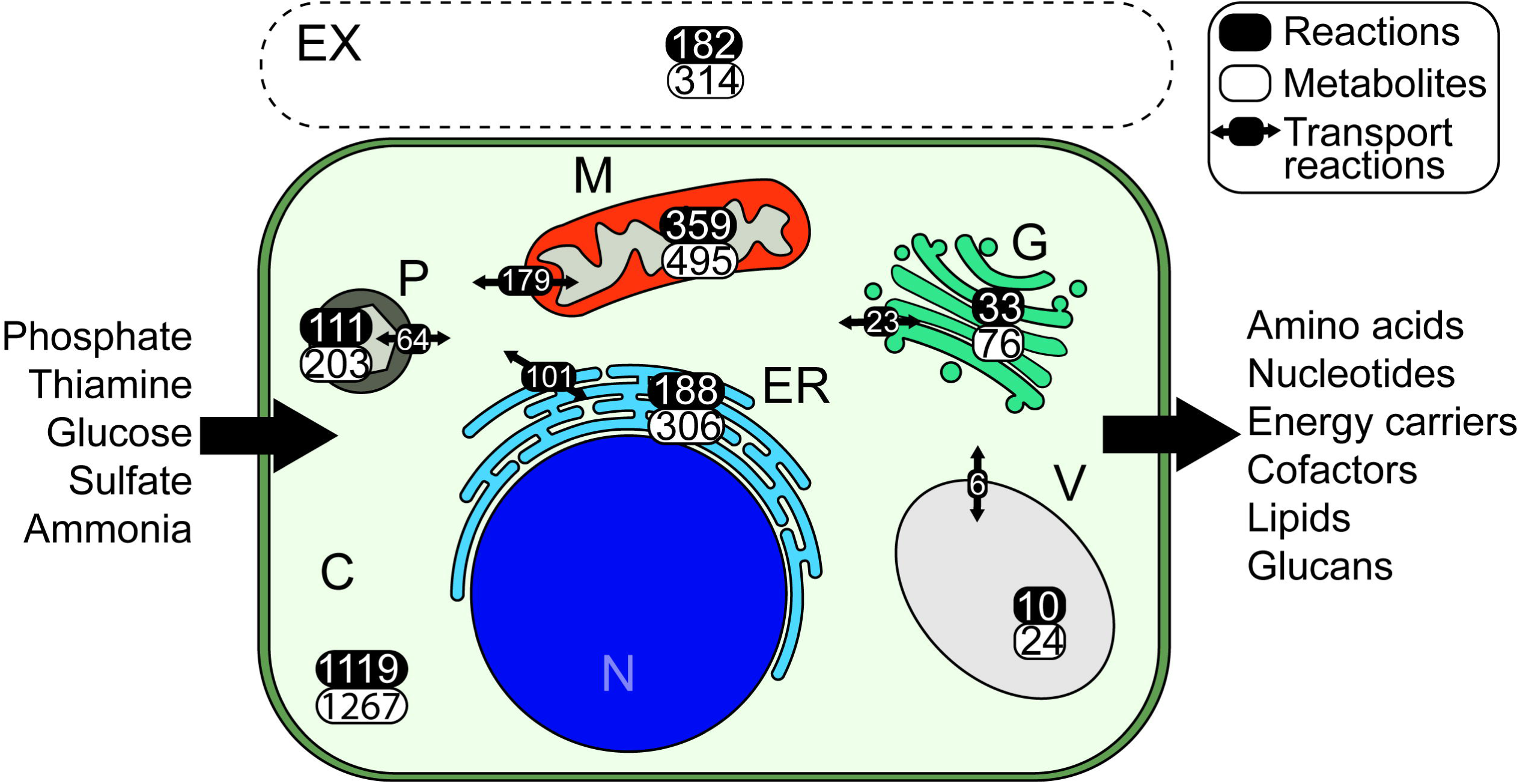
Schematic representation of a *Phytophthora infestans* cell with the number of reactions and metabolites per subcellular compartment and the number of transport reactions deduced from the model presented in this study. In this model the nucleus (N) is not included as a separate subcellular compartment. C: cytosol, M: mitochondrion, P: peroxisome, G: Golgi complex, V: vacuole, EX: extracellular space, ER: endoplasmic reticulum.

**Table 1:**
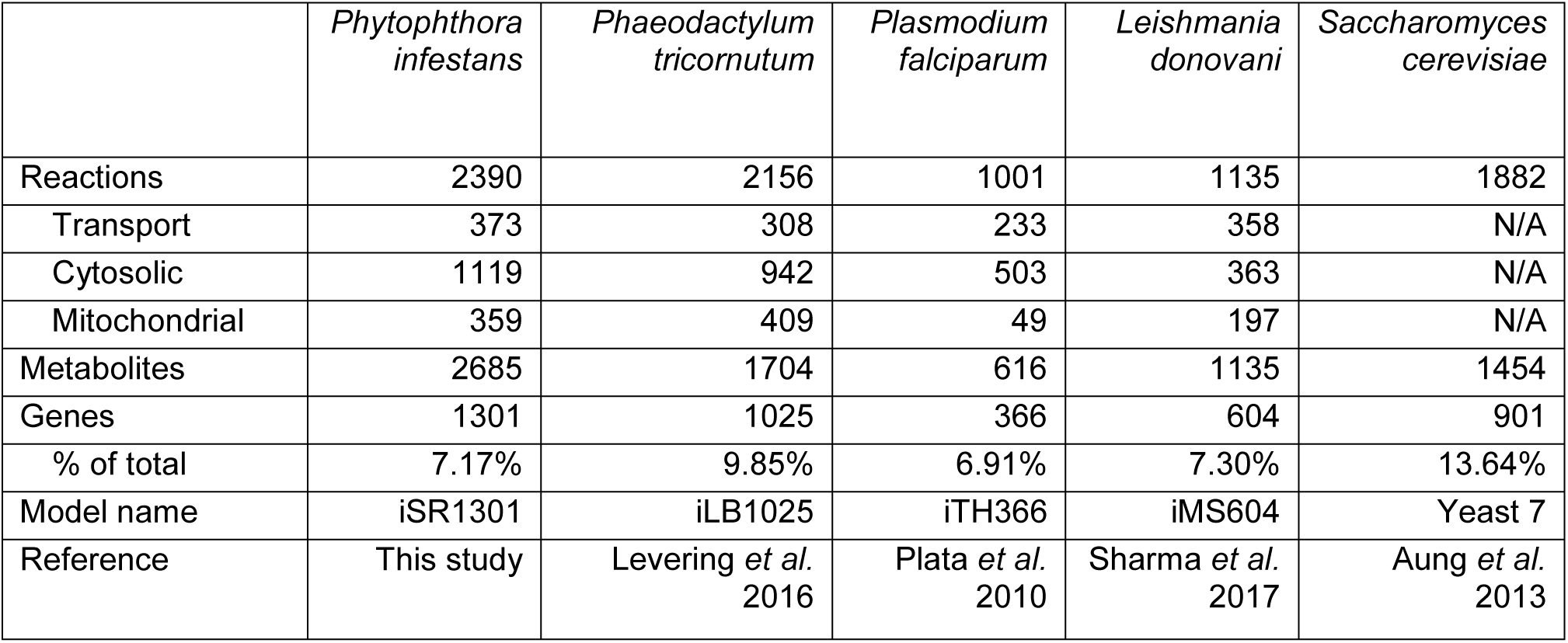
Statistics of the *Phytophthora infestans* GEM *iSR1301* and GEMs of other eukaryotic microbes.

In our model, the TCA cycle shares five reactions with the cytosol. One of these is catalyzed by malate dehydrogenase (MDH, EC 1.1.1.37). *P. infestans* has two genes encoding MDH, one encoding an isoform of MDH shown to be active in mitochondria in *P. infestans* and the other encoding a cytoplasmic isoform (López-Calcagno *et al.*, 2009). Other mitochondrial reactions are part of various metabolic pathways, including fatty acid biosynthesis, fatty acid degradation (β-oxidation) and even three glycolytic reactions, involving seven enzymes. The mitochondrial localization of these latter enzymes is likely a remnant of a secondary endosymbiosis event (Judelson, 2017).

### Model correction enables flux simulations

After initial reconstruction of the model, 153 invalid reactions (e.g. polymer reactions, see Methods) and 107 associated genes were removed from the model. The reconstructed metabolic model of *P. infestans* was initially unable to simulate growth (flux towards all biomass components), due to missing reactions or invalid reaction directionality constraints. This can be the result of an incomplete genome sequence or missannotations. We therefore performed a model gap-filling optimization to find the minimal set of reactions in KEGG that must be added to the model to correct this (Supplementary Table S2). This method proposed 16 additional reactions, and highlighted three reactions that must be reversed to allow for production of all biomass precursors. Notably, no gap-filling solutions were found for production of the fatty acids eicosapentaenoic acid (EPA) and behenic acid (AKA docosanoate), both of which are produced by *P. infestans* (Griffiths *et al.*, 2003; Robinson and Bostock, 2015; Sun *et al.*, 2013). This is caused by the lack of fatty acid reactions in KEGG, leaving multiple fatty acid reactions unconnected to other reactions (see KEGG map 01040).

To simulate the metabolite exchange between subcellular compartments, the model must also include intracellular transport reactions. Although nutrient transporters in *P. infestans* have been studied (Abrahamian *et al.*, 2016; Grenville-Briggs *et al.*, 2010), hardly anything is known about the metabolites that are exchanged between the cytosol and subcellular compartments. The annotated substrates for transporter proteins are not specific, and are therefore hard to integrate into the metabolic network. Moreover, transporter substrates such as those from the Transporter Classification Database (Saier *et al.*, 2016) are not cross linked with other databases. To overcome these limitations, we performed an optimization to identify the most likely set of intracellular transport reactions to be added to the network to allow production of all possible metabolites. We determined what metabolites could ultimately be produced by the model, after which we selected the set of transport reactions between the cytosol and any compartment to allow for this (Figure 1). The extracellular space (regarded as a subcellular compartment) was excluded from this optimization. The metabolism in this compartment is largely governed by cell wall degrading enzymes. Since it is not possible to distinguish the origin of the metabolites, the pathogen or the host plant, we had to exclude the extracellular space in this analysis.

After all correction steps, 1,290 of the 2,390 (53%) reactions in the model are able to carry flux based on the defined growth medium, 275 of which carry a nonzero flux when we calculate the optimal fluxes for maximal biomass production (Supplementary Table S1). Of the 2,685 metabolites in the model, 406 could not be produced based on our defined growth medium, and may require additional nutrient uptake. By iteratively adding uptake reactions to the model for each of these metabolites, we can simulate if the import of a specific metabolite would allow the production of additional metabolites (Supplementary Table S2). This reveals unresolved gaps in the model that could have a technical cause, but may also hint at biological properties. For example, episterol is proposed as a compound that would enable production of four other metabolites. This is striking since *Phytophthora spp.* lack sterol biosynthesis enzymes and depend on sterol acquisition from the host plant (Dahlin *et al.*, 2017). Another proposed metabolite is tyramine that would, upon import in the model, enable the production of six other metabolites. Tyramine is a product of decarboxylation of tyrosine, and based on the genome annotation *P. infestans* seems to lack the enzyme that catalyzes this reaction i.e. tyrosine decarboxylase (EC 4.1.1.25). However, a more precise examination of the genome sequence revealed an unannotated open reading frame (on supercontig 1.18, position 2365580 to 2367055) that likely encodes this enzyme.

### The metabolic model connects genomic and metabolic properties

We compared the properties of the *P. infestans* GEM to GEMs of other eukaryotic microbes (Table 1). The size of our *P. infestans* model, in terms of integrated reactions and genes, is in the same order of magnitude as that of a recent GEM of *Phaeodactylum tricornutum*, a closely related diatom (Levering *et al.*, 2016), although our model involves more metabolites. The sizes of the GEMs of the malaria parasite *Plasmodium falciparum* (Plata *et al.*, 2010) and the Leishmaniasis parasite *Leishmania donovani* (Sharma *et al.*, 2017) is much smaller, but the proportion of genes in the model is similar to that of the *P. infestans* model (~7% of total gene amount). Although these numbers might be smaller because of genome annotation quality and the level of model curation, it could also be due to the loss of primary metabolic pathways, for which these parasites rely on nutrient import from their hosts (Dean *et al.*, 2014; Gardner *et al.*, 2002). Despite the fact that *P. infestans* has a similar parasitic lifestyle, a pattern of pathway loss is not reflected in the size of our model.

The relation of a gene to an enzyme and its associated reactions is called the gene-protein-reaction (GPR) association (Machado *et al.*, 2016; Thiele and Palsson, 2010). A reaction can be associated to multiple enzymes (isozymes) and genes (paralogs). Conversely, one enzyme may present multiple catalytic domains, or it can have a broad substrate specificity, which associates it to multiple reactions. This “many-to-many-to-many” relationship holds information about redundancy of enzyme encoding genes in a genome, but also about gene essentiality, and the metabolic robustness of an organism to perturbations and different nutrient composition (Belda *et al.*, 2012). In our model, 40.4% of the genes are associated with just a single reaction, and 45.0% of the reactions in the model are associated with a single gene, which makes the respective genes essential for specific metabolic tasks (Supplementary Figure S2). In comparison, for the *P. tricornutum* GEM these numbers are higher (68.6% and 54.5% respectively). The diatom model is presumably of higher quality, since most reactions are manually curated. However, it might also hint at less redundancy of metabolic enzymes.

### Stage-specific models reflect reduced metabolic activity in sporangia and zoospores

It has been demonstrated that integration of transcriptomics data into a metabolic model has the potential to unveil condition- or tissue specific metabolic activity (Agren *et al.*, 2012; Becker and Palsson, 2008; Gatto *et al.*, 2014; Huthmacher *et al.*, 2010). We had access to transcriptome data of four asexual life stages, i.e. mycelium, sporangia, zoospores and germinating cysts (Schoina *et al.*, unpublished), and deployed the iMAT algorithm (Shlomi *et al.*, 2008) to predict stage-specific metabolic models for these life stages. This algorithm considers binary gene expression, i.e. a gene can either be expressed or not. Subsequently, it finds the fluxes through the model, supported by the maximum number of expressed genes, independent of defined medium and biomass composition. This results in sub-models for which all included reactions can carry flux. However, not all underlying genes have to be expressed. In other words, the resulting stage-specific models are sets of reactions that correlate best to the expression of the underlying genes. These reactions are therefore most likely to be metabolically active. If a reaction is absent from a stage-specific model, it is either absent because the expression of the associated genes is low, or because upstream reactions are absent. Comparing the sets of reactions in each stage-specific model, might reveal highly active life-stage specific metabolic activity. The distribution of stage-wise expression values for the genes in the model form a slimmer distribution (with slightly higher mean) than that of the total set of genes, indicating that genes in the model are more uniformly expressed (Supplementary Figure 3a). To generate a sufficiently large contrast between the stage-specific models we set the binary gene expression threshold at 7.04 TPM, the median of all expression values. Based on this threshold, genes were called expressed/not expressed, and life stage-specific models were calculated. Fewer genes were considered expressed in the sporangium and zoospore stages than in mycelium and germinating cyst (Supplementary Figure 3b).

The stage-specific models for sporangium and zoospore stages contain fewer reactions in total (Figure 2a), concordant with the observed general downregulation of many metabolic pathways in these stages (Ah-Fong *et al.*, 2017). The mycelium and germinating cysts models contain 907 and 905 reactions, respectively. Of these 891 are shared, indicating that these models are highly similar. Although the majority of the reactions, i.e. a core set of 795 reactions, is shared between all four stage-specific models there are also obvious differences; 57 reactions are specifically absent from the zoospore model (hence present in the other three), 19 reactions are only absent from the sporangium model while 20 reactions are absent from the sporangium and zoospore models, but present in mycelium and germinating cyst models. A principal component analysis on stage-wise reaction presence/absence (Figure 2b) shows that the mycelium and germinating cyst models cluster relatively close, whereas the sporangium and the zoospore models are more isolated. In summary, our metabolic flux data reflects the regulatory changes that reroute the metabolism of *P. infestans* during each life stage, especially the transitions between mycelium/germinating cyst and sporangium/zoospore stages (Ah-Fong *et al.*, 2017).

**Figure 2:**
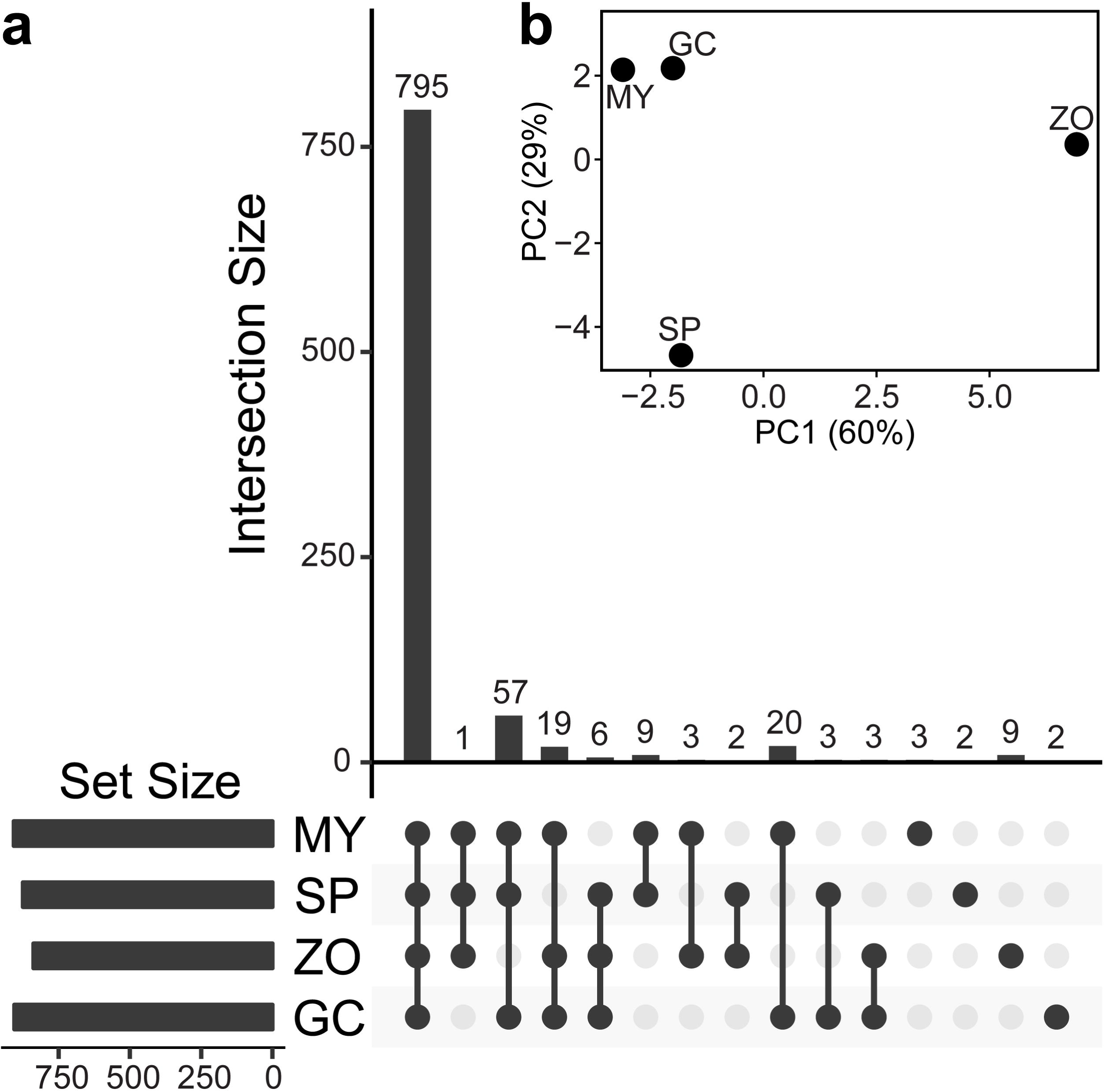
Stage-specific models of *Phytophthora infestans* mycelium (MY), sporangia (SP), zoospores (ZO) and germinating cysts (GC) a) Overlap of reaction content between the four stage-specific models. The bars on the bottom-left show the total numbers of reactions in each stage-specific model. The connected bullets indicate the models that are compared, and the bars in the graph represent the number of reactions (intersection size, Y-axis) that overlap between the stage-specific models. b) PCA on the stage-wise presence/absence (1/0) of a reaction in the stage-specific models.

To further interpret the presence/absence of reactions in the stage-specific models, we looked at the associated metabolic pathways (Supplementary Figure S4). For instance, the mycelium model contains two unique reactions of the “Vitamin B6 metabolism” pathway (KEGG R00173 and R00174), which represent the interconversion of pyridoxal (vitamin B6) to pyridoxal phosphate, an important cofactor for a large number of reactions, especially for the synthesis of amino acids (Percudani and Peracchi, 2003). The nitrogen metabolism pathway is represented by a core set of eight reactions, but the zoospore model lacks three reactions compared to mycelium and sporangia. Interestingly, these are reactions that contribute to glutamine and glutamate synthesis. Recently, the upregulation of nitrate transporters in zoospores has been reported (Ah-Fong *et al.*, 2017), which suggests an active nitrogen flux during this life stage. However, a reduced concentration of all amino acids was found in zoospores compared to other life stages (Grenville-Briggs *et al.*, 2005). As pointed out earlier the expression of enzymes in the nitrogen metabolism pathway is highly dynamic and depends on available nutrients (Abrahamian *et al.*, 2016). Possibly the nitrogen imported during the zoospore stage is stored and converted to amino acids at later life stages.

We observe the largest contrast of stage-wise reaction presence/absence in the fatty acid biosynthesis (FAB) pathway (Figure 3). A set of 13 reactions is present in the mycelium and germinating cyst models, and absent in the sporangium and zoospore models. Eight reactions are specifically absent in the zoospore model, but here six other reactions are specifically present. The latter are all mediated by two cytosolic fatty acid synthases (PITG_10922 and PITG_10926), seemingly downregulated in other stages. Instead, the mitochondrial fatty acid synthase (PITG_18025) seems active in the mycelium and germinating cyst stages. It is likely that fatty acids are synthesized during hyphal stages, since zoospores are thought to use stored fatty acids as nutrient source (Grant *et al.*, 1988; Yousef *et al.*, 2012). This data emphasize that fatty acids likely have an important role in *Phytophthora* zoospores. The three fatty acid synthase enzymes in *P. infestans* play a major role in the FAB process. Intriguingly, there could be a switch between cytosolic and mitochondrial FAB in zoospores. An unanticipated finding, both reported by Ah-Fong and colleagues (2017) and based on our model, is that fatty acid degradation (β-oxidation) is not pronounced in the zoospore stage, despite the predicted role of fatty acids in zoospore motility (Judelson, 2017).

**Figure 3:**
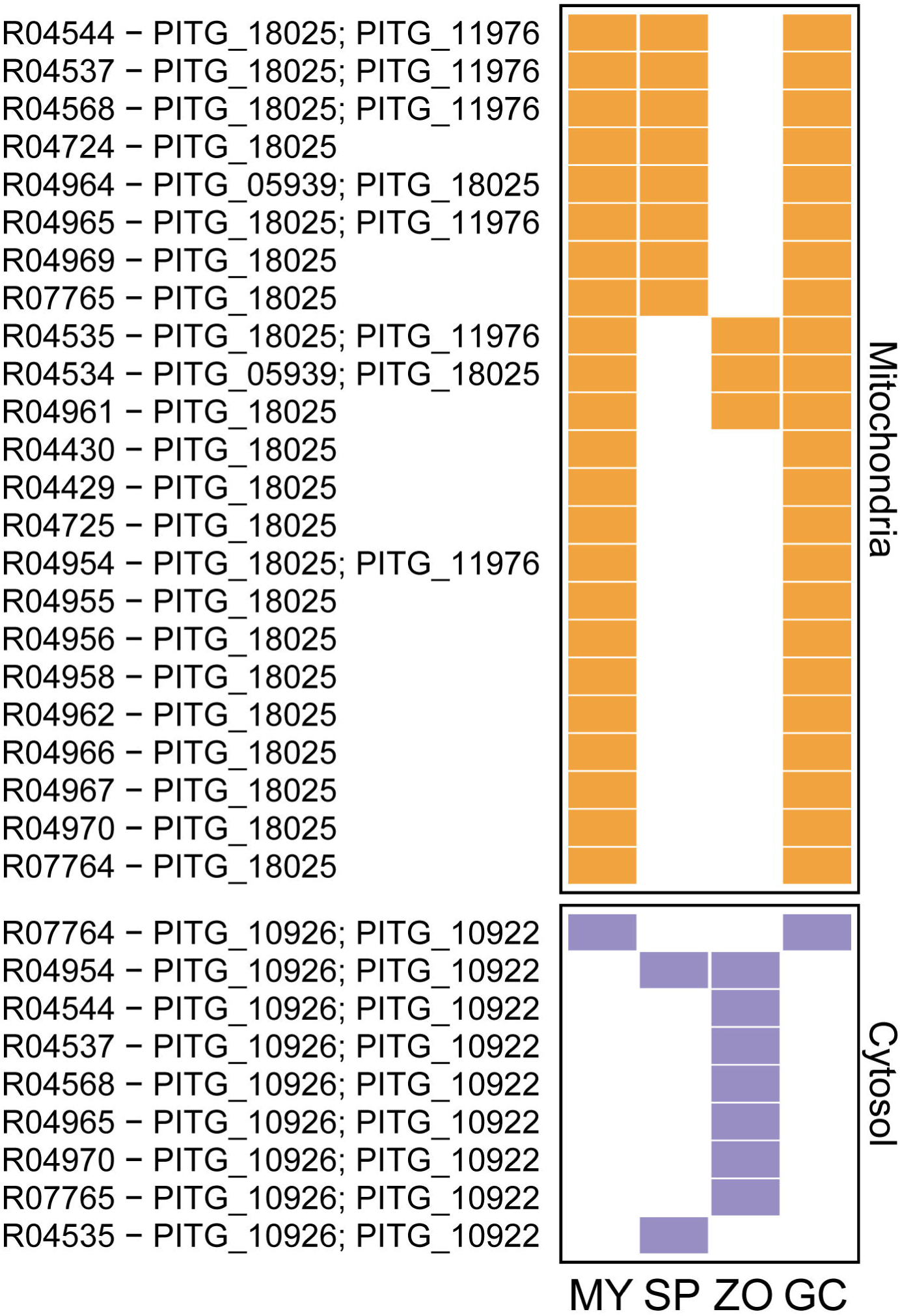
Fatty acid biosynthesis reactions in the stage-specific models of *Phytophthora infestans* mycelium (MY), sporangia (SP), zoospores (ZO) and germinating cysts (GC). Presence/absence of a KEGG reaction (indicated by its ID followed by associated gene IDs) is shown by filled/empty tiles, respectively while mitochondrial and cytosolic reactions are shown in orange and purple, respectively.

### Gene deletion simulations propose metabolic vulnerabilities

We investigated what effect gene deletions could have on the primary metabolism of *P. infestans.* By removing single genes from the model, one or more of the associated reactions in the model may be disabled. If such reactions are essential for production of any of the biomass precursors, these deletions disable growth i.e. the mathematical solution of the model becomes infeasible (O’Brien *et al.*, 2015), making such genes interesting candidates for further study. We performed single gene deletion (SGD) simulations of all genes in the model, which suggested 50 genes that would disable growth by disabling production of one of the essential biomass precursors (Supplementary File S2). These genes were associated with 163 reactions in various metabolic pathways (Figure 4). The pathways “phenylalanine, tyrosine and tryptophan biosynthesis” (17 out of 26 reactions vulnerable to SGD) and “valine, leucine and isoleucine” (11/17) are by far the most vulnerable pathways. Notably, the fatty acid degradation pathway is also delicate (17/50). In contrast, the most robust pathways are “tyrosine metabolism” (1/39), and “amino sugar and nucleotide sugar metabolism” (1/27).

**Figure 4:**
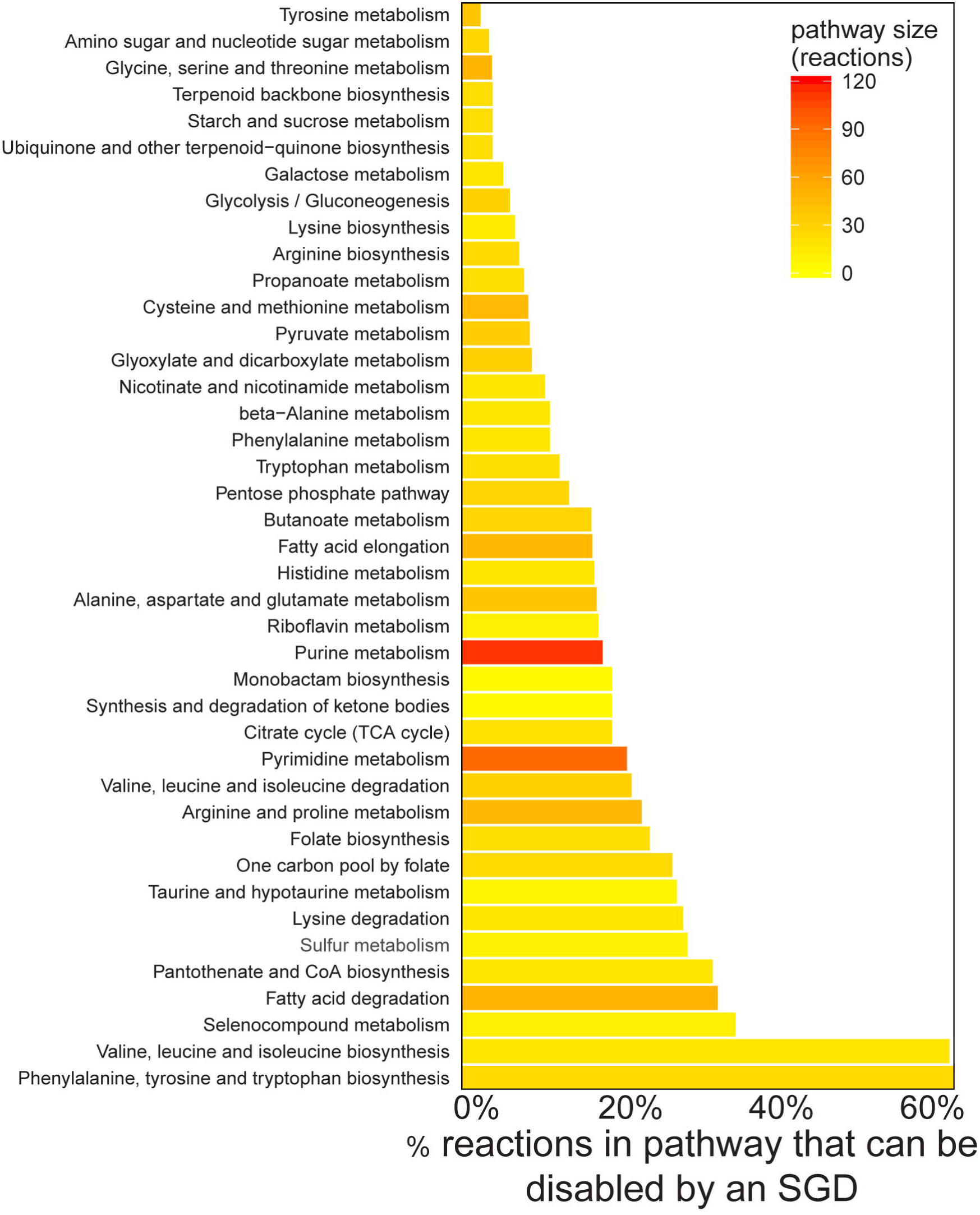
Gene deletion simulations in the *Phytophthora infestans* model. Percentage of reactions in each KEGG pathway that can be knocked out by a single gene deletion (SGD), disabling production of at least one biomass precursor. The colors of the bars scale with the absolute numbers of reactions found in our model in a particular KEGG pathway (inset top right).

There are numerous examples of the application of this method to suggest drug targets in pathogens (Hartman *et al.*, 2014; Kaltdorf *et al.*, 2016; Plata *et al.*, 2010; Sharma *et al.*, 2017; Yizhak *et al.*, 2013), thus suggesting that identified enzymes in *P. infestans* represent interesting candidates for further study. This analysis is of course based on *in vitro* growth conditions, while in its natural habitat *P. infestans* mainly resides *in planta.* It is thought that *P. infestans* imports larger, organic nutrients such as amino acids during infection (Abrahamian *et al.*, 2016).

## Concluding remarks

Here we present, to our knowledge, the first genome-scale metabolic model for the oomycete *Phytophthora infestans*, reconstructed mostly *in silico*, based on reactions found in KEGG. The aim of this study was not to provide a fully quantitative model, but rather to provide a broad overview of cellular metabolism, related to its genome. We optimized the model to be able to convert a minimal pool of nutrients into a set of minimal biomass precursors established from literature. Our model contributes to an understanding of the metabolism of *P. infestans.* However, even after gap filling, the fatty acids EPA and behenic acid could not be generated from the model, due to missing reactions in KEGG. This underscores the limitations of using solely KEGG as a resource, which may be overcome by manual refinement of fatty acid biosynthesis pathways from different databases, such as MetaCyc or BRENDA (Caspi *et al.*, 2014; Placzek *et al.*, 2017). In fact, the BRENDA database holds information on an omega-3 desaturase able to convert arachidonic acid into EPA (EC 1.14.19.25). This exact enzyme of *P. infestans* was recently proven capable of catalyzing this reaction in yeast (Yilmaz *et al.*, 2017). We also revealed one unannotated tyrosine decarboxylase enzyme in the *P. infestans* genome. These are clearly shortcomings that must be addressed in future versions of this model. Additionally, an experimentally assessed biomass composition, transporter integration, inclusion of species-specific reaction constraints and growth rates may be used to improve the accuracy of this model.

The life stage-specific models provide a direct functional context for the transcriptome data, by predicting the behavior of *P. infestans* metabolism under influence of stage-wise gene expression and are an alternative for the enrichment methods typically employed in metabolic pathway analyses. Approaching transcriptomic data from a functional point of view may emphasize certain features that are otherwise easily overlooked (e.g. the fatty acid biosynthesis). By building this model we can identify genes that have an essential role when converting simple nutrients to the building blocks of life. The metabolic model we reconstructed provides a scaffold for future genome-wide systems biology approaches to characterize the metabolism of *P. infestans*, and is an essential first step towards an integrative model of *P. infestans*–host interactions.

## Methods

### Draft reconstruction

We reimplemented the *getModelFromKEGG* method from the RAVEN Toolbox (Agren *et al.*, 2013) to improve performance and to incorporate minor adaptations. Briefly, Hidden Markov Models (HMMs) were trained on orthologous enzyme sequences derived from the KEGG Orthology (KO) database, Release 2015-11-23. We constructed multiple sequence alignments (MSA) of all eukaryotic protein sequences in every KEGG orthologous group using MAFFT version 7.273 (Katoh and Standley, 2013), using the “localpair” mode for local alignments. For performance reasons, the number of sequences used in an MSA was capped at 100, in which case we selected a random subset of sequences in the KO group. If fewer than 20 eukaryotic sequences were included in a KO group we constructed MSAs for prokaryotic sequences in respective KO group, maintaining the same rules. We used *hmmbuild* from the HMMER package version 3.1b (Eddy, 1998) to train HMMs on the MSAs. By using *hmmsearch*, we matched the HMMs to the protein sequences of *P. infestans* strain T30-4 (Haas *et al.*, 2009), at the time downloaded from the BROAD Institute website (https://www.broadinstitute.org), currently hosted at NCBI (bioproject 17665). Default parameters were maintained for hmmbuild and hmmsearch, and an E-value threshold of 10^−20^ was applied for hmmsearch. Similar to the *getKEGGModelForOrganism* function of the RAVEN Toolbox (Agren *et al.*, 2013), we performed two pruning steps, but we applied slightly stricter thresholds. First, any protein match to a KO group was removed if 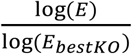 < 0.9, where *E* represents the *E*-value of respective protein to a KO group, and *E_bestKO_* represents the E-value of the best-matching KO group for that protein. In other words, protein hits are often removed if they have a better match to another KO group. Second, any protein match to a KO group was removed if 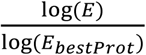 < 0.5, where *E* again represents the E-value of respective protein match to a KO group, and *E_bestProt_* represents the lowest E-value of any protein to this KO group. This reduces the number of matches per KO group to reduce the number of false positives, since there is clearly a better matching protein. Subsequently, we retrieved all KO annotations of *P. infestans* from KEGG (organism ID “pif”). The combined set of matched KO groups was used to retrieve all associated reactions and metabolites from KEGG. Consequently, each reaction in the model was associated to a number of *P. infestans* genes. We hypothesized that each of the genes associated to a reaction is able of catalyzing it. We did not consider enzyme complexes, which together would fulfil a single enzymatic task. Reactions were removed automatically if their stoichiometry was undefined (e.g. “1,3-beta-D-Glucan(n) + UDP-D-glucose <=> 1,3-beta-D-Glucan(n+1) + UDP”), and if the same metabolite ID occurred at both sides of the reaction arrow, which implies a polymer reaction (e.g. “UTP + RNA <=> Diphosphate + RNA”). Additionally, reactions that were associated to metabolites containing the substrings “acceptor”, “donor”, “tRNA”, “enzyme”, “aglycon” and “fatty acid” were removed.

### Model correction

We predicted the subcellular location of *P. infestans* proteins using LocTree 3 (Goldberg *et al.*, 2014), and subsequently distributed the associated reactions of the model over the cellular compartments. We selected seven compartments for our model: cytosol, extracellular space, mitochondria, endoplasmic reticulum, Golgi complex, peroxisome and vacuole. Reactions associated to enzymes with transmembrane predictions (plasma or intracellular membranes) were assigned to respective compartments on both sides of the membrane, since it is unclear where the catalytic domain is localized. Proteins assigned to any other than our seven compartments, were assigned to the cytosol. Next, all proteins that were predicted to be secreted by Raffaele and colleagues (2010) were assigned to the extracellular space compartment.

The initially reconstructed model was exported to SBML and Microsoft Excel format using CobraPy v0.4.1 (Ebrahim *et al.*, 2013). For the next steps we imported the model into MATLAB (R2015b) using the RAVEN Toolbox v1.8 (Agren *et al.*, 2013). We used Gurobi v7.0.1 (http://www.gurobi.com/) to solve the (mixed-integer) linear programs.

Nutrient uptake and biomass reactions were added to the model by applying the *fillGaps* function from the RAVEN Toolbox to propose gap solutions from KEGG, by constraining biomass production to a positive flux. This method implements the SMILEY algorithm (Rolfsson *et al.*, 2011), including all reactions from a universal set of reactions (in this case KEGG), and subsequently minimizing the flux through these. Prior to this, we temporarily removed the extracellular space compartment to prevent gap solutions here, and added all possible transport and excretion reactions to the model. To predict the set of transport reactions between compartments, we assessed which metabolites can ultimately be produced by the model. Then, we constrained a positive flux for the production of these metabolites, and we removed the transport reactions that did not carry flux.

After these correction steps, we used the function *solveLP* from the RAVEN Toolbox to solve the linear programs of flux balance analysis (FBA), and to get the flux distribution for maximal biomass production. We applied the function *checkProduction* to detect metabolite gaps in the model. This method checks which metabolites are not producible (blocked) from the model, and then iteratively adds uptake reactions for these metabolites, to see whether the uptake of these metabolites unblocks metabolites elsewhere in the model.

### Stage-specific models

We used RNA sequencing data (Schoina *et al.*, unpublished) to quantify gene expression in four *in vitro* life stages of *P. infestans*, i.e. mycelium, sporangia, zoospores and germinating cysts. Gene expression of *P. infestans* strain T30-4 (Haas *et al.*, 2009) was quantified using Kallisto v0.42.4 (Bray *et al.*, 2016), which expresses mRNA abundance in transcripts-per-million (TPM). This unit represents the number of reads aligned to transcript sequences, normalized for transcript length and sequencing depth, scaled by a million. We determined a binary expression threshold to define if a gene is expressed or not, for which we used the median of all expression values over the four life stages. We decomposed the biomass and other excreted compounds (lipids etc.) into separate excretion reactions, and used the iMAT algorithm (implemented in the function *createTissueSpecificModel)* incorporated in the COBRA Toolbox (Schellenberger *et al.*, 2011; Shlomi *et al.*, 2008) to derive stage-specific models.

### Gene knockout simulations

We used the function *findGeneDeletions* from the RAVEN Toolbox to predict the genes that would, upon deletion, disable biomass flux. This method first selects reactions that are supported by a single gene, and then iteratively constrains the flux of these reactions to zero. A gene is marked as essential if the linear program of FBA becomes infeasible, i.e. when no flux through the biomass reaction is possible.

## Acknowledgements

This work was partly supported by the Food-for-Thought campaign from the Wageningen University Fund and by The Netherlands Organization for Scientific Research in the framework of a VENI grant (M.F.S.).

## Supporting information

**Supplementary Figure S1:** Reactions in the *Phytophthora infestans* model per subcellular compartment and overlap in reaction content between different subcellular compartments. These include cytosol (cyto), mitochondrion (mito), extracellular space (extr), endoplasmic reticulum (ER), peroxisome (pero), Golgi complex (golg), and vacuole (vacu). The bars on the bottom-left show the total numbers of reactions in each subcellular compartment. The connected bullets indicate the compartments that are compared, and the bars in the graph represent the number of reactions (intersection size, Y-axis) that overlap between the compartments.

**Supplementary Figure S2:** Frequencies of (a) gene numbers per reaction and (b) reaction numbers per gene in the *Phytophthora infestans* model.

**Supplementary Figure S3:** Transcriptome data of *Phytophthora infestans* in relation to stage-specific metabolic models. a) Distributions of transcripts-per-million (TPM) expression values of all *P. infestans* genes (background) and the genes in the model, combined from four life stages. The red line indicates the TPM threshold set to distinguish expressed/non-expressed genes in the model. b) The percentages of all *P. infestans* genes (background), and the genes in the model for which gene expression in each life stage exceeds the TPM threshold. MY: mycelium; SP: sporangia; ZO: zoospores; GC: germinating cysts.

**Supplementary Figure S4:** Numbers of reactions per KEGG pathway that are shared between the *Phytophthora infestans* life stage specific models of mycelium (MY), sporangia (SP), zoospores (ZO) and germinating cysts (GC). The colors of the tiles scale to the relative frequencies of all non-core reactions (i.e. the reactions that are absent in at least one stage-specific model). The numbers in the two right-most columns represent the core set of reactions (shared by all stage-specific models) and the total set of reactions for respective pathway (core + non-core).

**Supplementary Table S1:** Properties of all reactions in the *Phytophthora infestans* model.

**Supplementary Table S2:** Gap-filling solutions, candidate metabolites, and essential reactions in the *Phytophthora infestans* model.

**Supplementary SBML File S1:** The *Phytophthora infestans* model in SBML format.

